# Differential Effects of SUMO1/2 on Circadian Protein PER2 Stability and Function

**DOI:** 10.1101/789222

**Authors:** Ling-Chih Chen, Yung-Lin Hsieh, Tai-Yun Kuo, Yu-Chi Chou, Pang-Hung Hsu, Wendy W. Hwang-Verslues

## Abstract

Posttranslational modification (PTM) of core circadian clock proteins, including Period2 (PER2), is required for proper circadian regulation. PER2 function is regulated by casein kinase 1 (CK1)-mediated phosphorylation and ubiquitination but little is known about other PER2 PTMs or their interaction with PER2 phosphorylation. We found that PER2 can be SUMOylated by both SUMO1 and SUMO2; however, SUMO1 versus SUMO2 conjugation had different effects on PER2 turnover and transcriptional suppressor function. SUMO2 conjugation facilitated PER2-β-TrCP interaction leading to PER2 proteasomal degradation. In contrast, SUMO1 conjugation, mediated by E3 SUMO-protein ligase RanBP2, enhanced CK1-mediated PER2^S662^ phosphorylation and increased PER2 transcriptional suppressor function. PER2 K736 was critical for both SUMO1- and SUMO2-conjugation. A PER2^K736R^ mutation was sufficient to alter circadian periodicity and reduce PER2-mediated transcriptional suppression. Together, our data revealed SUMO1 versus SUMO2 conjugation acts as an upstream determinant of PER2 phosphorylation and thereby affects the circadian regulatory system and circadian periodicity.

## Introduction

The circadian clock controls many rhythmic physiological processes such as hormonal oscillation, metabolism and immune function that are essential to maintain homeostasis (Dunlap, 1999; Stephan & Zucker, 1972). At the molecular level, the circadian clock establishes these biological rhythms through interconnected transcription-translation feedback loops (TTFLs). The core loop is composed of *BMAL*, *CLOCK*, Period (*PER1*, *PER2*) and Cryptochrome (*CRY1*, *CRY2*). BMAL-CLOCK heterodimers transcriptionally activate the expression of PER and CRY and other circadian output genes. In turn, PER and CRY suppress their own expression and expression of other circadian output genes controlled by the BMAL-CLOCK complex (Hughes et al., 2009; Shearman et al., 2000). PER2 suppresses BMAL/CLOCK mediated transcription by displacing BMAL/CLOCK/CRY complexes from target promoters. This displacement is not only important for inhibition of circadian gene expression, but also essential for reactivation of the TTFL (Chiou et al., 2016). This core loop regulates expression of retinoic acid receptor–related orphan receptor (ROR) and REV-ERBα/β (NR1D1, NR1D2) which form a secondary loop that stabilizes the core loop by regulating BMAL1 and CRY1 transcription (Bugge et al., 2012; A. C. Liu et al., 2008; Preitner et al., 2002).

The robustness and stability of circadian rhythm is also reinforced by post-translational modifications (PTMs) that determine the level, activity and subcellular localization of core clock proteins (Gallego & Virshup, 2007; Hirano, Fu, & Ptacek, 2016; C. Lee, Etchegaray, Cagampang, Loudon, & Reppert, 2001; Partch, Green, & Takahashi, 2014; Reischl & Kramer, 2011; Stojkovic, Wing, & Cermakian, 2014). Although the core clock proteins are known to be regulated by several types of PTMs, it is likely that additional layers of clock-related PTM regulation remain to be discovered. Among the known PER2 PTMs, phosphorylation at two important serine residues (S662, S480) has been most intensively studied (C. Lee et al., 2001). Phosphorylation at S662 affects PER2 subcellular localization and therefore influences transcription of PER2 itself and its downstream genes (Vanselow et al., 2006; Y. Xu et al., 2007). Phospho-null mutation of PER2 S662 enhances PER2 repressor function but increases PER2 turnover in the nucleus. Loss of PER2 S662 phosphorylation (typically the result of a PER2 S662G substitution) causes human Familial Advanced Sleep Phase Syndrome (FASPS) which is characterized by a dramatically shortened circadian period with a 4-hour advance of sleep, temperature, and melatonin rhythms (Toh et al., 2001; Y. Xu et al., 2007). Phosphorylation at S480 facilitates PER2-β-TrCP interaction leading to PER2 proteasomal degradation (β-TrCP phosphodegron) (Eide et al., 2005). Acetylation of PER2 at lysine (K) residues also protects PER2 from ubiquitination; whereas deacetylation of PER2 by SIRT1 leads to degradation (Asher et al., 2008).

Recent research has shown that a phosphoswitch where CK1δ/ε can phosphorylate either of the two competing PER2 phosphorylation sites, S662 and S480, determines PER2 stability (Fustin et al., 2018; Narasimamurthy et al., 2018; Zhou, Kim, Eng, Forger, & Virshup, 2015). However, the factors that control this switch and determine which site CK1 will phosphorylate remain unknown. This illustrates how, even for a relatively well studied protein like PER2, interaction and cross regulation between PTMs that control protein function can be complex and not well understood. It is known that PER2 phosphorylation can be influenced by other PTMs. For example, O-GlcNAcylation in a cluster of serines around S662 competes with phosphorylation to regulate PER2 repressor activity (Kaasik et al., 2013). Whether other PTMs are also part of this interactive regulation of PER2 that influences the speed of circadian clock (Masuda et al., 2020) remains unknown.

Despite these many lines of evidence showing the important of PER2 PTMs, little is known about PER2 SUMOylation. SUMOylation can alter protein-protein interactions, subcellular localization or protein activity and can participate in transcriptional regulation (Gareau & Lima, 2010; Gill, 2005). Usually, SUMOylation does not promote protein degradation. However, for a subset of proteins, conjugation with multiple SUMOs is critical for recognition by SUMO-targeted ubiquitin ligases (STUbLs), leading to proteasomal degradation (Hunter & Sun, 2008). At least three major SUMO proteins have been identified in higher eukaryotes: SUMO1 and SUMO2/3. SUMO2 and SUMO3 are 97% identical, but they share less than 50% sequence identity with SUMO1. SUMO2/3 often form SUMO chains on the substrates, whereas SUMO1 typically appears as monomers or acts as a chain terminator on SUMO-2/3 polymers (Matic et al., 2008). However, SUMO1 and SUMO2/3 may still be able to act redundantly as SUMOylation events which typically use SUMO1 can be compensated by SUMO2 or SUMO3 in SUMO1-deficient mice (Evdokimov, Sharma, Lockett, Lualdi, & Kuehn, 2008). Conversely, conjugation by SUMO1 or SUMO2/3 can impart different fates to the SUMOylated protein. SUMO1 conjugation typically affects cellular processes including nuclear transport, cell cycle control and response to virus infection (Saitoh, Pu, & Dasso, 1997), antagonizes ubiquitination (Desterro, Rodriguez, & Hay, 1998) and modulates protein-protein interaction (Gill, 2004). In contrast, SUMO2/3 is thought to mainly participate in cellular response to stress (Saitoh & Hinchey, 2000) and can facilitate targeted protein ubiquitin-mediated degradation (Tatham et al., 2008). Recently, SUMO1 and SUMO2/3 have been found to have opposite functions in mouse lens cells at different stages of development by differentially regulating the transcriptional activity of Specificity Protein 1 (SP1) (Gong et al., 2014). Whether or not other transcriptional regulators are similarly affected by SUMO1 versus SUMO2/3 conjugation is unknown.

We found that PER2 K736 is critical for both SUMO1 and SUMO2 conjugation. However, SUMO1 versus SUMO2 conjugation had dramatically different effects on PER2 protein turnover and transcriptional suppressor function. PER2 SUMO1 conjugation acted upstream of and was critical for CK1-mediated S662 phosphorylation and PER2 transcriptional repressor activity. In contrast, SUMO2 promoted PER2 degradation. These data show that SUMOylation is an important layer of PER2 regulation which impacts the critical phosphoswitch controlling PER2 function and coordination of the circadian regulatory system.

## Results

### PER2 can be SUMOylated

Serum shock to synchronize and induce circadian gene expression (Balsalobre, Damiola, & Schibler, 1998) showed that PER2 had a peak and trough of expression at circadian times (CT) of 28 and 40 h, respectively, in U2OS cells (Fig. 1A). Immunoprecipitation (IP) of PER2 at these times of maximum and minimum expression found multiple PER2-SUMO species (Fig. 1B). This indicated that PER2 was either SUMOylated at multiple sites or conjugated to multi-SUMO chains. Moreover, the molecular weight of SUMOylated PER2 differed between CT28 and CT40. Stronger PER2-SUMO1 signal at lower molecular weight close to the unmodified PER2 was detected at CT28 (indicated by *); whereas this signal was not apparent at CT40 (Fig. 1B). Proximity ligation assay (PLA) showed that SUMO1-PER2 interaction was mainly detected in the nucleus at CT28 (Fig. 1C). However, the interaction signals was also found in the cytoplasm in more than half of the cells examined at CT40 and the portion of cells with nuclear only or nuclear and cytoplasmic PER2-SUMO1 differed between different CT times (Fig. 1C). The different SUMO1-PER2 species observed and subcellular localization at different CT indicated that SUMOylation may contribute to PER2 oscillation or circadian function.

**Figure 1.**
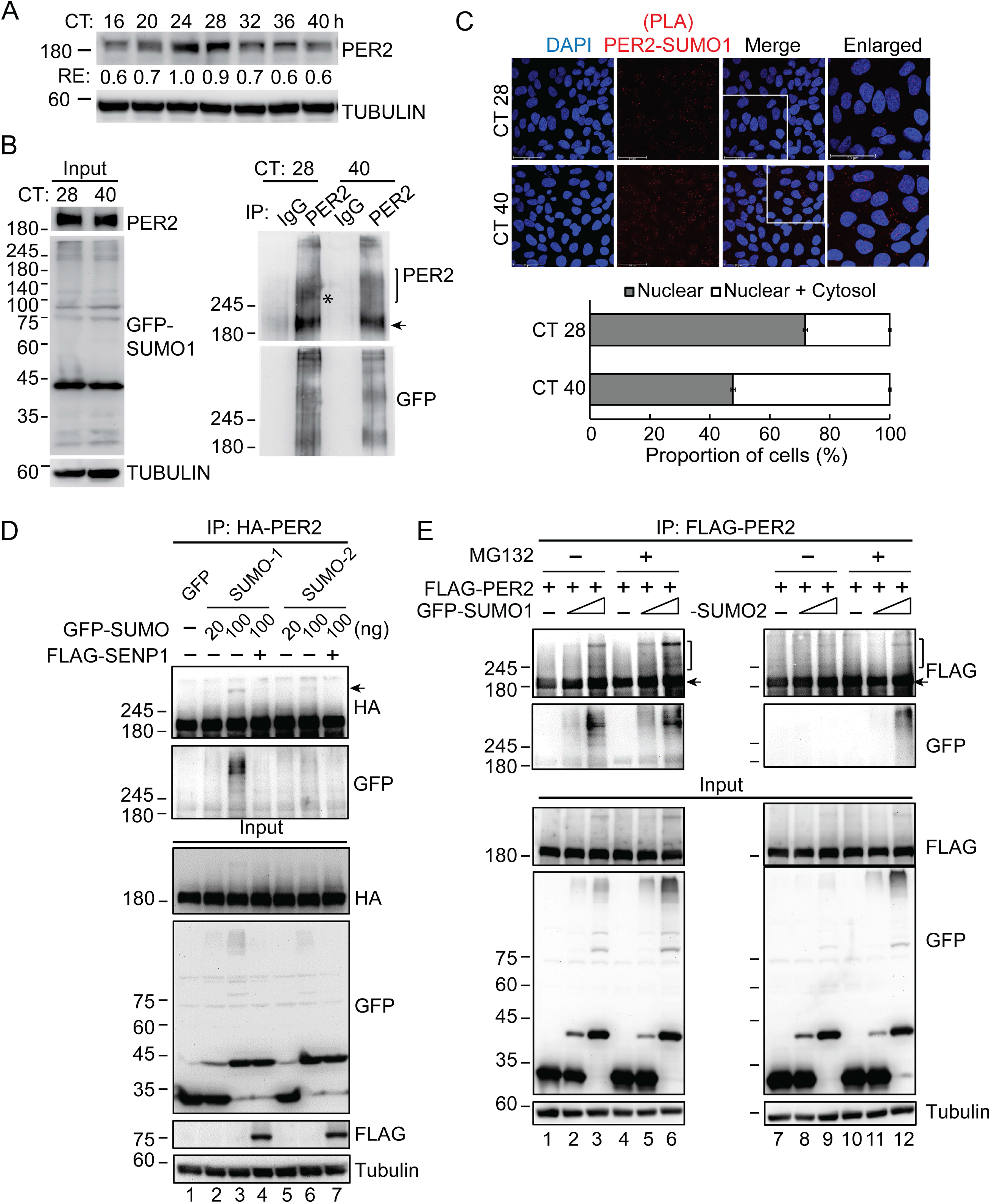
Differential SUMOylation of PER2 is detected at different circadian times. **A.** Time course assay using synchronized U2OS cells. After 50% serum shock, total protein was extracted at the indicated circadian time (CT). The level of endogenous PER2 was determined using immunoblotting (IB) analysis. Tubulin was used as a loading control. RE: relative expression. **B.** Co-immunoprecipitation (Co-IP) assay using GFP-SUMO1 expressing U2OS cells at CT28 and CT40 after serum shock. Cell lysates were immunoprecipitated with anti-PER2 antibody. Immunoprecipitates and input lysates were analyzed by IB. Arrow indicates the unmodified PER2 and brackets indicated PER2-SUMO1 and PER2-multi-SUMOs conjugates. Asterisk (*) indicates the stronger PER2-SUMO1 signal at lower molecular weight close to the unmodified PER2. The experiment was repeated twice. **C.** Proximity ligation assay (PLA) using U2OS cells at CT28 and CT40 after serum shock. Cells were fixed and PLA was performed using anti-PER2 and anti-SUMO1 antibodies against endogenous PER2 and SUMO1 proteins. Top: Representative confocal microscopy images are shown for each condition, which was repeated at least three times. Scale bar, 50 μm. Bottom: The proportion of different subcellular localization patterns of PER2-SUMO1 interaction was calculated from nine non-overlapping confocal images for each CT (326 and 508 cells were counted for CT28 and CT40, respectively). Data are proportion of cell numbers ± 95% confidence intervals. **D.** Co-IP assay using HEK-293T cells co-transfected with HA-PER2, GFP-SUMOs and FLAG-SENP1. Cell lysates were IP with anti-HA antibody. Immunoprecipitates and input lysates were analyzed by IB. Arrow indicates the PER2-SUMO conjugates. **E.** Co-IP assay using HEK-293T cells co-transfected with FLAG-PER2 and GFP-SUMO1 or GFP-SUMO2. Cells were also treated with the proteasome inhibitor MG132. Cell lysates were IP with anti-FLAG antibody and analyzed by IB. Arrows indicate the PER2-SUMO conjugates.

PER2-SUMOylation was further confirmed by detection of SUMO-conjugated PER2 in HEK-293T cells ectopically expressing HA-taggedPER2 and GFP-SUMO1 or GFP-SUMO2 (Fig 1D, lane 3 and 6). Co-expression of SUMO1 or SUMO2 with SENP1, a SUMO protease, eliminated the PER2 band shift (Fig 1D, lane 4 and 7), confirming that PER2 could be SUMOylated by both SUMO1 and SUMO2. While these data indicated that PER2 could be conjugated to both SUMO1 and SUMO2, the signal of PER2-SUMO2 modification was less prominent than that of PER2-SUMO1 (Fig 1D, lane 6 vs. lane 3). One possible explanation is that SUMO2 modification leads to PER2 ubiquitination and proteasomal degradation. Consistent with this hypothesis, treatment with the proteasome inhibitor MG132 resulted in accumulation of PER2-SUMO2 conjugates (Fig 1E, lane 12 vs. lane 9).

### SUMO2 promotes PER2 protein degradation

Further experiments validated our hypothesis that SUMO2 conjugation promotes PER2 degradation. PER2 protein levels were unaffected by SUMO1 but decreased in a dose-dependent manner upon SUMO2 expression (Fig 2A; S1A). Immunoblot (IB) assay using whole cell lysates from cycloheximide (CHX)-treated cells further showed that SUMO2, but not SUMO1, significantly shortened PER2 protein half-life (Fig 2B; S1B). To demonstrate that SUMO2-conjugation promoted PER2 degradation via ubiquitination, Co-IP assay was performed using HEK-293T cells ectopically expressing FLAG-PER2 and GFP-SUMO2. Addition of MG132 to inhibit proteasomal degradation resulted in accumulation of not only PER2-SUMO2 conjugates, but also ubiquitinated-PER2 species (Fig S1C, lane 5, 6 vs. lane 2, 3).

**Figure 2.**
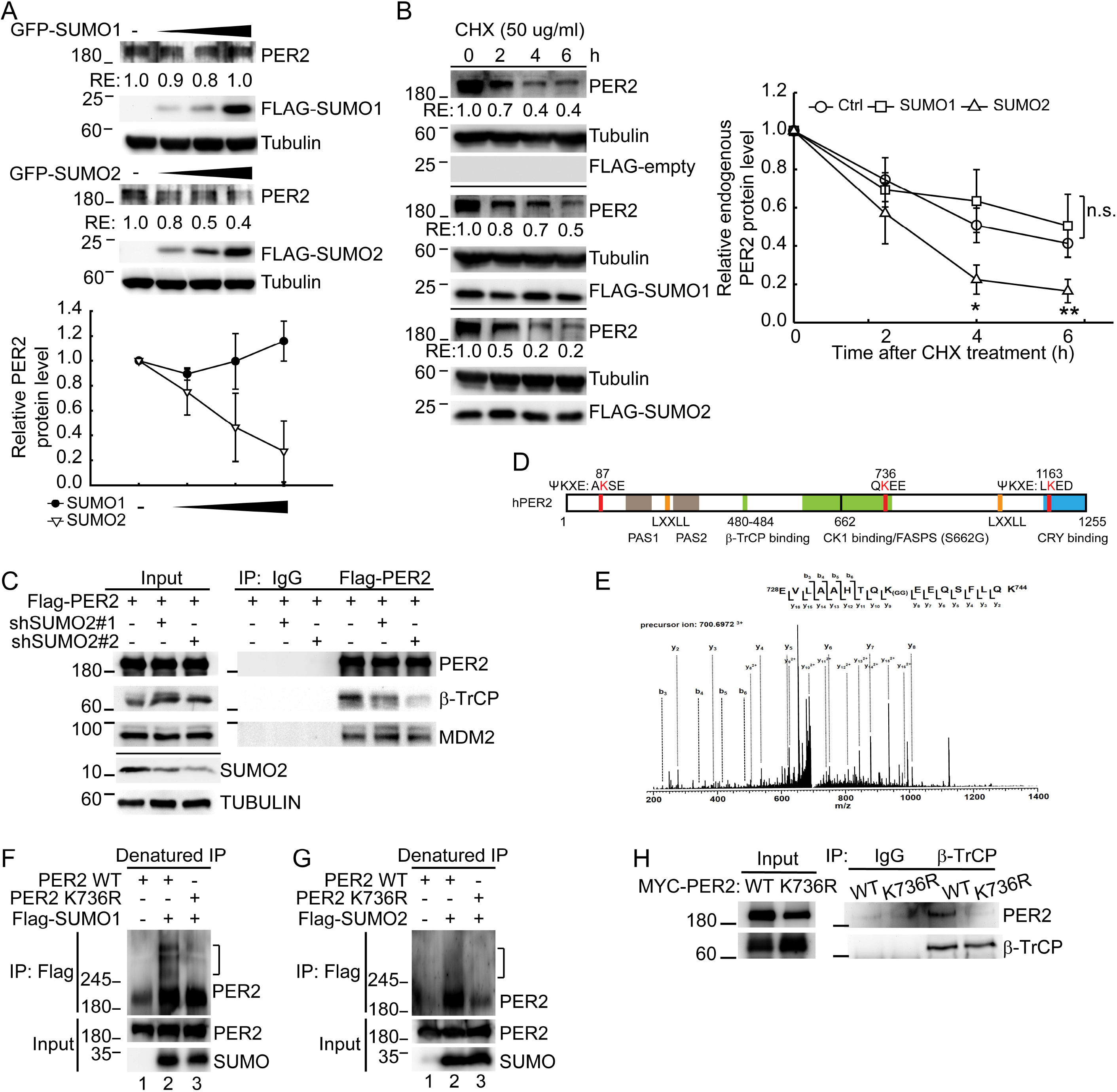
Lysine (K) 736 is important for PER2 SUMOylation and SUMO2 conjugation on K736 facilitates PER2 protein degradation via promotion of PER2-βTrCP interaction. **A.** IB assay of endogenous PER2 and FLAG-SUMOs using U2OS cells transfected with increasing amount of FLAG-SUMOs. Tubulin was used as a loading control. RE: relative expression. Relative PER2 protein levels from three IB analyses were quantified. Data are means ± SD. **B.** Time course assay using cycloheximide (CHX) treated U2OS cells transfected with FLAG-SUMOs or FLAG-empty control. The levels of endogenous PER2 and FLAG-SUMOs were determined using IB analysis. Tubulin was used as a loading control. RE: relative expression. Right panel: Quantification of relative endogenous PER2 protein levels from six IB analyses. Data are means ± SD. n.s., non-significant; *, *p* < 0.05; **, *p* < 0.01 (Student’s t-test). **C.** Co-IP assay of PER2 and β-TrCP or MDM2 using U2OS cells transduced with shSUMO2 lentiviral vectors. Cell lysates were IP with indicated antibodies and analyzed by IB assay. Normal IgG was used as an IP control. The experiment was repeated twice. **D.** Schematic representation of human PER2 protein. Three predicted SUMOylation sites (K87, K736, K1163) are indicated (red). Coactivator LXXLL nuclear receptor recognition motifs (orange), CK1 targeted serine regions (S480, S662, green), the familial advanced sleep phase syndrome (FASPS) mutation site (S662G, black), β-TrCP binding, Per-Arnt-Sim (PAS) domains (brown) and a CRY binding domain (blue) are also indicated. **E.** The MS/MS spectrum of the tryptic peptide m/z 700.6972 from PER2 with SUMOylated K736.All b and y product ions from the SUMOylated peptide with di-glycine modification on K736 are labelled in the spectrum. The experiment was repeated twice. **F.** Denatured IP of PER2 and SUMO1 using HEK-293T cells co-transfected with MYC-PER2 and FLAG-SUMOs. Cell lysates were IP with anti-FLAG antibody and analyzed by IB analysis. Brackets indicate the PER2-SUMO conjugates. The experiment was repeated twice. **G.** Denatured IP of PER2 and SUMO2 using HEK-293T cells co-transfected with MYC-PER2 and FLAG-SUMOs. Replication and data presentation are as described for E. **H.** Co-IP assay of PER2 and β-TrCP using HEK-293T cells transfected with MYC-PER2^WT^ or – PER2^K736R^. Cell lysates were IP with indicated antibodies and analyzed by IB. Normal IgG was used as an IP control.

PER2 can be degraded via S480 phosphodegron recognized by β-TrCP (Eide et al., 2005; Reischl et al., 2007) as well as phosphorylation-independent interaction with MDM2 (J. Liu et al., 2018). Therefore we performed Co-IP assay using SUMO2-depleted U2OS cells to determine which ubiquitin E3 ligase is responsible for SUMO2-mediated PER2 degradation (Fig. 2C). Upon SUMO2 depletion, PER2-β-TrCP interaction was decreased; whereas the interaction between PER2 and MDM2 was not affected (Fig. 2C) indicating that SUMO2-conjugation is important for β-TrCP-mediated PER2 degradation.

### Lysine 736 is important for PER2 SUMOylation

The majority of known SUMO substrates are SUMOylated at a lysine in the consensus motif *ψ*KxD/E (where *ψ* is a large hydrophobic residue) (J. Xu et al., 2008). However, SUMOylation can also occur at lysine residues outside this motif. Two putative PER2 SUMOylation sites at K87 and K1163 were identified using the GPS-SUMO bioinformatics database (Zhao et al., 2014) (Fig 2D). An additional SUMOylation motif, QKEE (Hendriks et al., 2017), was also identified at K736 by manual inspection of the PER2 sequence (Fig 2D).

To determine which of these putative PER2 SUMOylation sites are functionally important, a series of K to Arginine (R) mutations at K87, K736 and/or K1163 were created. The K736R mutation clearly inhibited SUMO2-mediated PER2 degradation while K87R and K1163R had no effect (Fig S2A). We further tested PER2 double and triple mutants and found that whenever K736 was mutated, PER2 protein became more stable regardless of whether K87 or K1163 could be SUMOylated (Fig S2B). These data indicated that K736 could be an important SUMOylated site of PER2. Consistent with this, liquid chromatography–tandem mass spectrometry (LC-MS/MS) using lysates prepared from HEK-293T cells transiently expressing FLAG–PER2 and EGFP‐SUMO1 identified a SUMO peptide branch at K736 of PER2 (Fig 2E). In addition, denatured-IP assays confirmed that the PER2 K736R mutation significantly decreased both SUMO1 and SUMO2 conjugation (Fig 2F, 2G). Moreover, the PER2 K736R mutation eliminated the band shift (Fig. S2C, lane 4 vs. lane 3) that was at the same position as the eliminated shift resulting from co-expression of SUMO1 with SENP1 (Fig S2C, lane 2 vs. lane 3). This further confirmed that PER2 could be SUMOylated at K736. Also consistent with the critical importance of PER2 K736, Co-IP showed a significant decrease in PER2^K736R^ interaction with β-TrCP (Fig 2H). While the mass spectrometry data demonstrated that K736 is itself SUMOylated, we do not rule out the possibility that mutation of K736 also affects SUMOylation at other unidentified sites. Together these results demonstrated that PER2 K736 is critical for PER2 SUMOylation.

### PER2 K736 is critical for maintenance of correct PER2 oscillation and circadian period

Stable clones of U2OS cells harboring the PER2^K736R^ mutation were generated by CRISPR/CAS editing (Fig S3). This endogenous PER2^K736R^ mutant was more stable and did not show rhythmic oscillation after serum shock compared to the PER2^WT^ (Fig. 3A, 3B). PER2^WT^ protein reached peak levels at CT24 and CT28 and reduced to the trough levels at CT16 and CT40. In contrast, PER2^K736R^ protein level remained at constant level throughout the time course (Fig. 3A, 3B). Analysis of circadian rhythm using a transiently expressed PER2 luciferase reporter showed that the PER2^K736R^ cells had significant lower circadian peak amplitude and shorter period compared to PER2^WT^ cells (Fig 3C). These observations demonstrated that PER2 K736, and K736-dependent SUMOylation, were required for proper circadian regulation.

**Figure 3.**
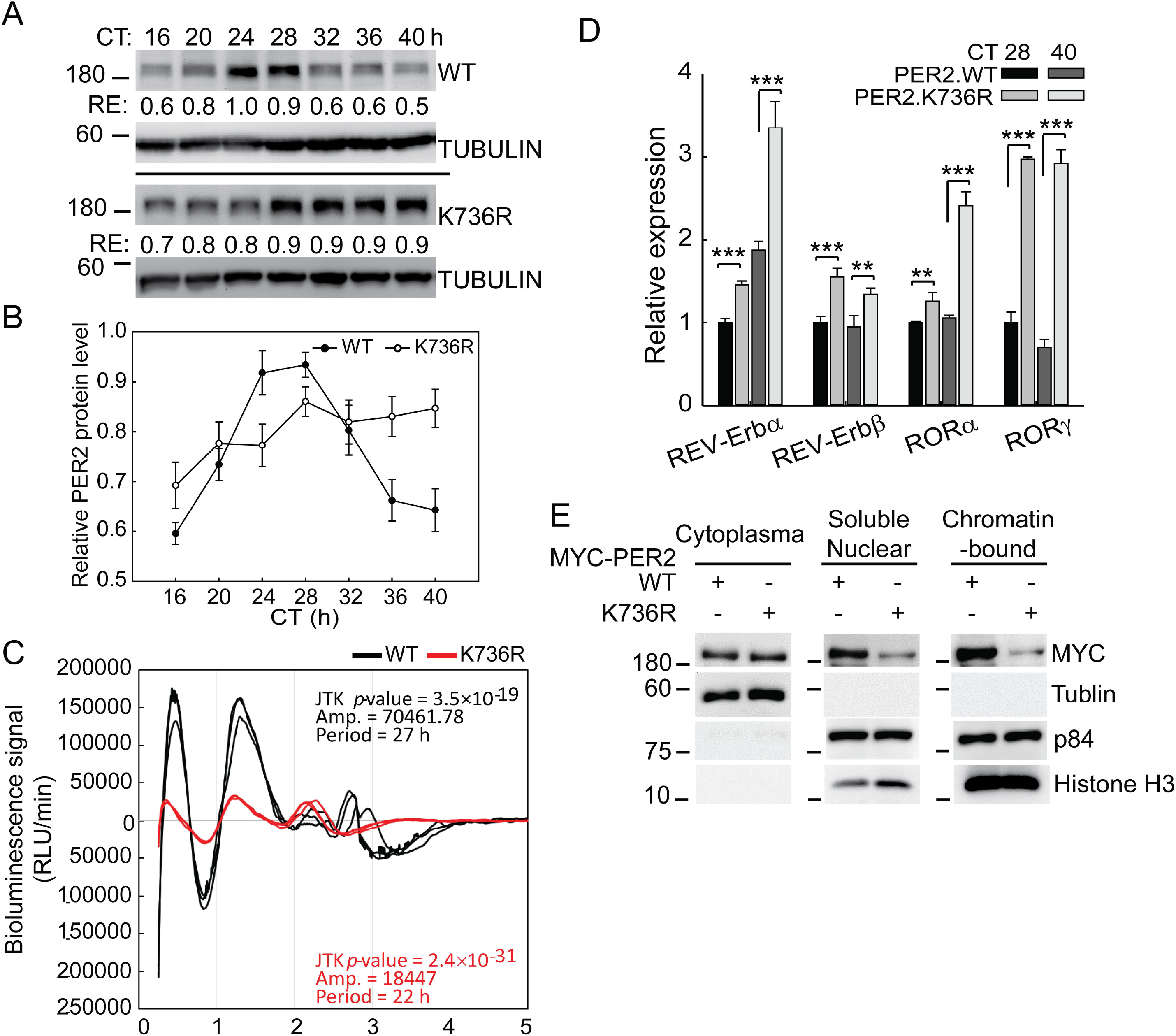
K736-dependent SUMO conjugation is required to maintain correct PER2 oscillation, PER2 nuclear localization, PER2-mediated transcriptional suppression and correct circadian period. **A.** Time course assay using synchronized U2OS cells with or without CRISPR/CAS-edited PER2^K736R^ knock-in. After 50% serum shock, total protein was extracted at the indicated circadian time (CT). The levels of endogenous PER2 and SUMOs were determined using IB analysis. Tubulin was used as a loading control. RE: relative expression. **B.** Quantification of relative PER2 protein levels from six time course analyses. Data are means ± SD. **C.** Circadian rhythm of bioluminescence using CRISPR/CAS-edited PER2^K736R^ knock-in or control (PER2^WT^) U2OS cells transiently transfected with PER2 luciferase reporter. Data were smoothed and detrended with the KronosDio analysis tool and the rhythmicity (JTK *p*-value), amplitude (Amp) and period were analyzed using JTK_Cycle analysis (version 3). n=3 **D.** Expression of genes in auxiliary circadian loops analyzed by real-time quantitative (q)-PCR using PER2^K736R^ knock-in or PER2^WT^ control U2OS cells at CT 28 and CT 40 after serum shock. GAPDH was used as an internal control. All data points were performed in at least triplicates and all experiments were performed at least twice with similar results. Data are presented as mean ± SD, n=3. **, *p* < 0.01; ***, *p* < 0.001 (Student’s t-test) **E.** Fractionation analysis using HEK-293T cells co-transfected with MYC-PER2^WT^ or −PER2 ^K736R^. The levels of MYC-PER2 in cytoplasmic, soluble nuclear and chromatin-bound fractions were determined using IB. Tubulin was used as a loading controls for cytoplasmic proteins. Nuclear matrix protein p84 and histone H3 were used as loading controls for nuclear proteins. The experiment was repeated twice.

### SUMOylation is critical for PER2 mediated transcriptional suppression

As PER2 is key player in the core circadian TTFL, changes in its subcellular localization and protein stability are expected to affect circadian-regulated gene expression, particularly genes in auxiliary feedback loops such as nuclear receptors REV-ERBs and RORs (Akashi & Takumi, 2005; Preitner et al., 2002). Therefore, we examined the expression of REV-ERBα, REV-ERBβ, RORα and RORγ in U2OS cells expressing PER2^WT^ or endogenous PER2^K736R^.

In PER2^WT^ cells, REV-ERBα, but not REV-ERBβ, RORγ or RORα, was de-repressed at CT40 when PER2^WT^ expression decreased to the trough level (Fig. 3A, 3B; 3D, dim gray vs. black). This result was consistent with the transcriptional suppressor function of PER2 in the TTFL. In contrast, REV-ERBα expression was higher in PER2^K736R^ mutant cells compared to the PER2^WT^ cells at CT28 (Fig. 3D, gray vs. black), and it reached an even higher level in PER2^K736R^ mutant cells at CT40 when the level of PER2^K736R^ remained high (Fig. 3A, 3B; 3D). The higher level of REV-ERBα at CT40 could be due to an increase in BMAL1 expression since BMAL1 and PER2 express in antiphase to each other. Also, PER2^K736R^ cells had higher expression of REV-ERBβ, RORγ and RORα than PER2^WT^ at both CT28 and CT40 (Fig 3D). These results confirmed that K736 modification was essential for PER2 transcriptional repression.

### PER2-SUMO1 conjugation promotes CK1 phosphorylation of PER2 S662 and GAPVD1 interaction required for PER2 nuclear translocation

PER2 must enter the nucleus to act as a transcriptional repressor. To determine if PER2 nuclear entry was affected by SUMOylation, we performed subcellular fractionation experiment using HEK-293T cells ectopically expressing MYC-tagged PER2^WT^ or PER2^K736R^ (Fig. 3E). A significantly lower amount of PER2^K736R^ was found in the soluble nuclear and chromatin-bound fractions compared to the PER2^WT^ (Fig. 3E). PER2 enters the nucleus as a complex with CK1 and CRY1 (Chaves et al., 2006; K. Miyazaki, Mesaki, & Ishida, 2001; Ollinger et al., 2014; Vanselow et al., 2006). Recent findings have also demonstrated that the interaction with GAPVD1, a Rab GTPase guanine nucleotide exchange factor (RAB-GEF) involved in endocytosis and cytoplasmic trafficking, is critical for the nuclear entry of the PER2/CRY/CK1 complex (Aryal et al., 2017). GAPVD1 had significantly decreased interaction with PER2^K736R^ (Fig S4A). Together, these results indicated that SUMO1-conjugation of PER2 facilitates PER2 nuclear entry by promoting PER2-GAPVD1 interaction and subsequent nuclear import of the PER2/CRY/CK1 complex.

Phosphorylation affects PER2 localization (Vanselow et al., 2006; Y. Xu et al., 2007) and rhythmic changes in PER2 phosphorylation are crucial for modulating circadian rhythm (Eide et al., 2005; Hirota et al., 2010; H. M. Lee et al., 2011; Shanware et al., 2011; Tsuchiya et al., 2009). Two functional CK1 phosphorylation sites have been identified in human PER2, S480 (S478 in mPer2) and S662 (S659 in mPer2) (Eide et al., 2005; Vanselow et al., 2006). Comparing our data with previous results, it was intriguing to note that PER2 K736-SUMO1 conjugation and S662 phosphorylation both led to increased PER2 nuclear retention (Fig 3E) (Vanselow et al., 2006). Conversely, K736-SUMO2 conjugation and S480 phosphorylation both promoted PER2 ubiquitination and proteasomal degradation (Fig 1E, 2A, S1A, S1B) (Eide et al., 2005; Reischl et al., 2007). Inhibition of CK1 using the CK1δ/ε inhibitor PF670462 led to reduced PER2 S662 phosphorylation but did not affect PER2 SUMOylation (Fig S4B), indicating that PER2 SUMOylation does not depend upon PER2 phosphorylation. Conversely, it has been shown that SUMOylation could induce phosphorylation (Luo et al., 2014). We therefore checked whether SUMOylation affects CK1-mediated PER2 phosphorylation and found that PER2^K736R^ was less efficiently phosphorylated by CK1 than PER2^WT^ (Fig S4C). This occurred despite the fact that PER2^WT^ and PER2^K736R^ had similar protein-protein interaction with CK1. Taken together, these data indicate that K736-mediated SUMO1 conjugation may serve as a signal to promote both CK1 phosphorylation of S662 and protein interactions needed for PER2 translocation into the nucleus.

### PER2 SUMO1 conjugation is mediated by RanBP2

Unlike ubiquitination, E3 ligases are not required for protein SUMOylation as the SUMO E2-conjugating enzyme UBC9 is able to conjugate SUMO onto target proteins (Melchior, 2000; Muller, Hoege, Pyrowolakis, & Jentsch, 2001). In the case of PER2, however, both SUMO1 and SUMO2 can be conjugated on the same K736 site. Therefore, a specific SUMO E3‐ligating enzyme may be required to mediate SUMO1 versus SUMO2 conjugation and thereby determine PER2 protein fate. As SUMO1 conjugation promoted PER2 nuclear entry, we hypothesized that the nucleoporin RanBP2, an E3 enzyme known to mediate SUMO1 conjugation to target proteins for nuclear import (Gareau & Lima, 2010; Pichler, Gast, Seeler, Dejean, & Melchior, 2002), could be responsible for PER2-SUMO1 conjugation. Reduction of PER2-SUMO1 conjugation was also observed in U2OS cells upon RanBP2 depletion (Fig. 4A). Consistent with this, depletion of RanBP2 in HEK-293T cells (Fig S4D) ectopically expressing MYC-PER2 and GFP-SUMO1 or GFP-SUMO2 resulted in a significant decrease in PER2-SUMO1 conjugation (Fig S4E, upper right panel). The effect of RanBP2 depletion on SUMO2 was less pronounced (Fig S4E, lower right panel). RanBP2 depletion significantly decreased PER2 S662-phosphorylation but did not affect interaction between PER2 and CK1 (Fig. 4B, 4C). This observation further demonstrated that PER SUMOylation occurs upstream of S662 phosphorylation. Together, these results indicate that RanBP2-mediated PER2-SUMO1 conjugation is critical for CK1-mediated PER2 phosphorylation at S662 to promote PER2 nuclear entry.

**Figure 4.**
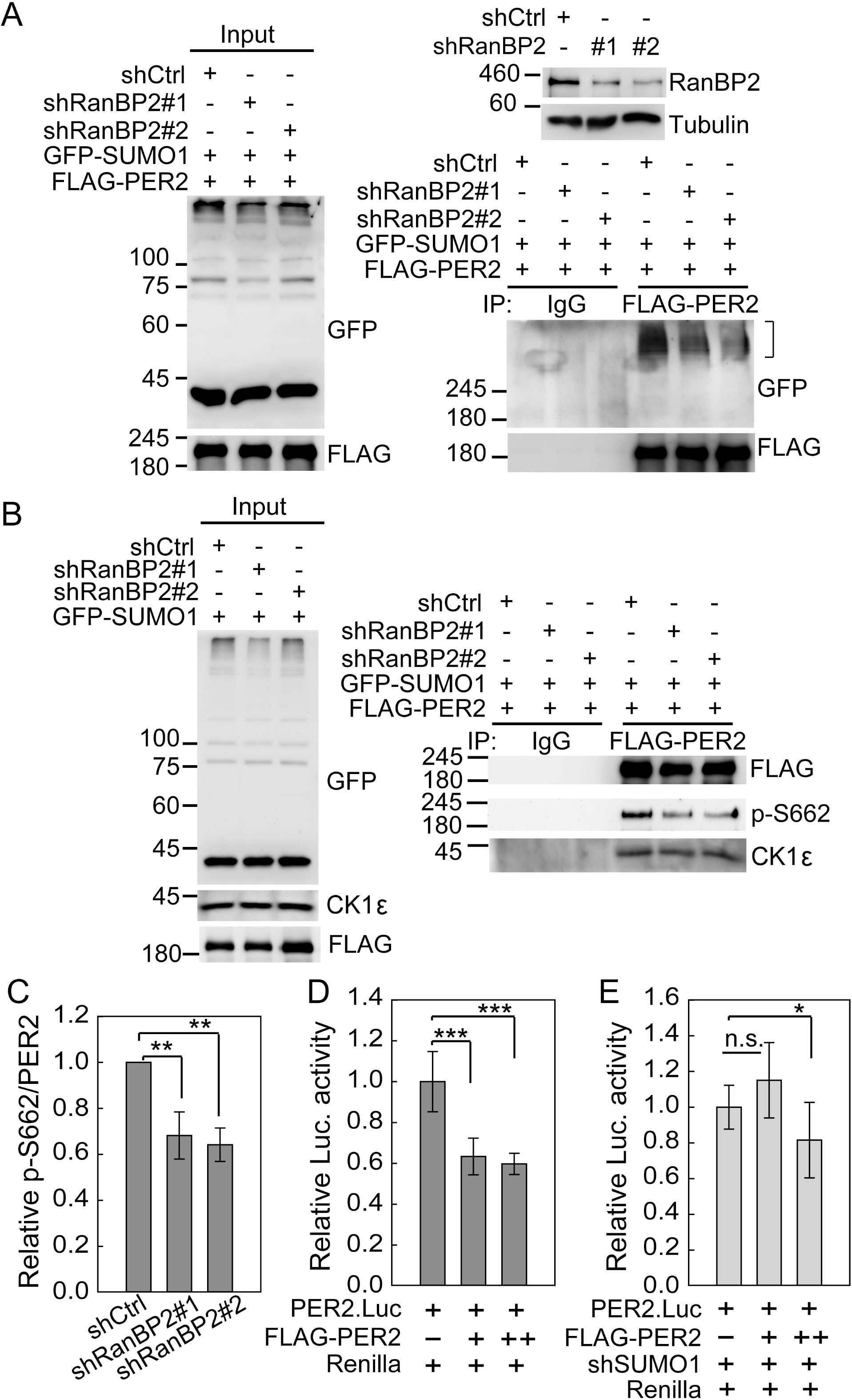
PER2 SUMO1 conjugation is mediated by RanBP2. **A.** Co-IP assay using U2OS cells without or with RanBP2 depletion co-transfected with FLAG-PER2 and GFP-SUMO1. Cell lysates were IP with anti-FLAG antibody and analyzed by IB to detect PER2-SUMO1 (lower right panel) conjugates indicated by brackets. The depletion efficiency of RanBP2 protein was determined using IB analysis (upper right panel). **B.** Co-IP assay using U2OS cells without or with RanBP2 depletion co-transfected with FLAG-PER2 and GFP-SUMO1. Cell lysates were IP with anti-FLAG antibody and analyzed by IB to detect CK1 and S662-phosphorylated PER2. **C.** Quantification of relative S662-phosphorylated PER2 protein levels (p-S662/FLAG-PER2) from three Co-IP analyses. Data are means ± SD. **D.** Relative fold-change in luciferase activity of the PER2 promoter-reporter construct in HEK-293T cells transiently co-transfected with FLAG-PER2^WT^. Luciferase activity in samples without FLAG-PER2^WT^ transfection was arbitrarily set at 1. Data are means ± SD, n=3. *, *p* < 0.05; ***, *p* < 0.001 (Student’s t-test). All data points were performed in at least triplicates and all experiments were performed at least two times with similar results. **E.** Relative fold-change in luciferase activity of the PER2 promoter reporter construct in HEK-293T cells transiently co-transfected with FLAG-PER2^WT^ with SUMO1 depletion. Replication and data analyses are as described for D.

**Figure 5.**
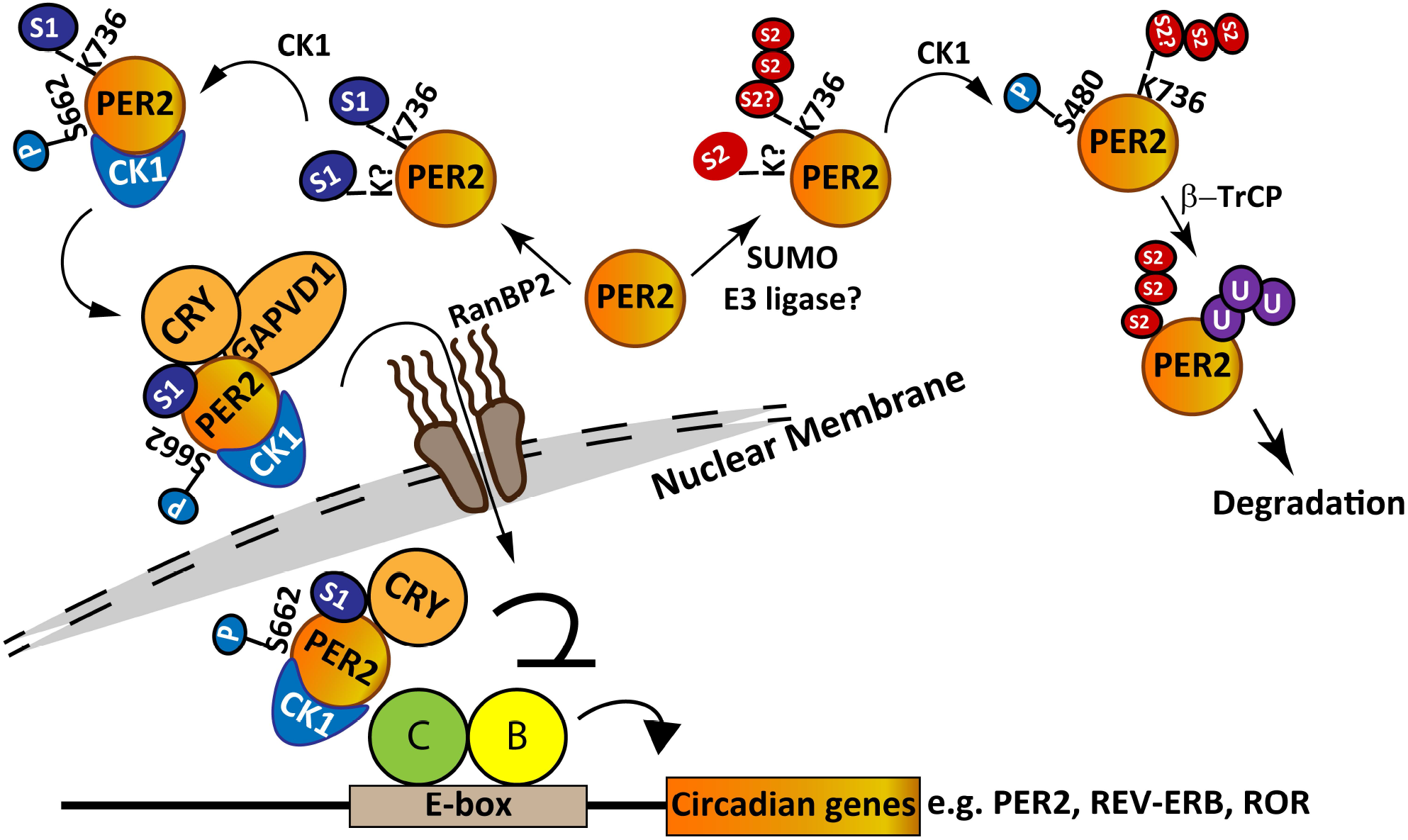
Diagram of the proposed mechanism by which SUMOylation regulates PER2 protein stability and transcriptional suppression function. PER2 K736 is important for PER2 SUMOylation by multiple SUMO isoforms. SUMO2 facilitates PER2 interaction with β-TrCP, ubiquitination and degradation. In contrast, SUMO1 modification does not affect PER2 protein stability, but enhances PER2 S662-phosphorylation and interaction with CRY1 and GAPVD1 resulting in an increase in nuclear PER2. Such modification is critical for PER2-mediated repression of circadian genes including PER2 itself and other genes such as REV-ERBα in auxiliary loops. The SUMO E3 ligase RanBP2 facilitates PER2-SUMO1 conjugation which is critical for CK1-mediated PER2 S662 phosphorylation. Identification of SUMO E3 ligases involved in SUMO2 modification of PER2 and analysis of whether SUMOylated K736 affects SUMOylation at other K sites is of interest for further experiments.

In the core TTFL, S662-phosphorylated PER2 dimerizes with CRY1 to suppress its own expression. To further demonstrate that SUMO1 is critical for PER2 transcriptional suppressor function, a transient reporter assay was performed. Expression of PER2^WT^ repressed PER2 promoter activity in HEK-293T cells (Fig 4D). However, such suppression was abolished upon SUMO1 depletion (Fig. 4E). These results were consistent with decreased PER2 nuclear import when SUMO1 expression was reduced and further demonstrated that K736 SUMOylation is required for PER2 to function as transcriptional suppressor regulating circadian genes.

## Discussion

Our data indicate that, compared to other well-established PER2 PTMs, SUMOylation has an equally large role in determining PER2 function and in maintaining correct circadian periodicity. Consistent with previous findings that S662 phosphorylation mainly promotes PER2 protein nuclear retention and stabilization (Vanselow et al., 2006), our data suggested that RanBP2-mediated PER2-SUMO1 conjugation enhances CK1-mediated PER2 phosphorylation at S662 and promotes PER2 transcriptional suppression function (Fig. 4). Conversely, PER2-SUMO2 conjugation, in line with CK1-mediated PER2 S480-phosphorylation and the β-TrCP-phosphodegron (Eide et al., 2005; Narasimamurthy et al., 2018), facilitated PER2-β-TrCP interaction, ubiquitination and degradation (Fig. 2). The circadian phosphoswitch of PER2 between S662 and S480 sites is determined by CK1 (Narasimamurthy et al., 2018). Our data showed that depletion of SUMO1 or K736R mutation did not affect CK1 binding to PER2 protein but reduced S662 phosphorylation (Fig.4, S4). This indicates that SUMO1 modification on K736 serves as a signal for CK1 to phosphorylate S662. Likewise, SUMO2 modification on K736 may prompt CK1 to phosphorylate S480. However, S480 is more distal to K736 than S662 and it is not known how SUMO2 conjugation at K736 may affect S480 phosphorylation (or vice versa). Taken together, our data indicate that PER2 SUMOylation is a primary PTM that controls downstream PTMs (phosphorylation and ubiquitination) to determine PER2 function and stability.

Recent evidence indicates that many SUMO targets can be modified by both SUMO1 and SUMO2/3 (Becker et al., 2013). However, the mechanisms balancing SUMO1 versus SUMO2/3 conjugation are unclear. One possibility is that differential SUMOylation is primarily due to the relative expression level of SUMO isoforms (Gong et al., 2014). Another possibility is that SUMO1 and SUMO2/3 may have different conjugation site preferences (Becker et al., 2013; Impens, Radoshevich, Cossart, & Ribet, 2014). In the case of PER2, K736 is important for both SUMO1 and SUMO2 conjugation (Fig 2). Whether PER2-SUMO2 conjugation requires a specific SUMO E3 ligase or can be carried out by the SUMO-conjugating enzyme UBC9 is of interest for future research.

It should also be mentioned that PER2 SUMOylation at K736 site is a unique regulation in primates. Sequence alignment from NCBI protein database showed that while primate PER2 sequence is QKEE in this region (amino acids 735/736 to 738/739) the corresponding region in rodents is QREE (Fig. S5). We demonstrated that expression of PER2^K736R^ resulted in a shorter circadian period in U2OS cells. Thus it will be interesting to further investigate whether this PER2 sequence divergence between primates and rodents is responsible for the shorter circadian period of rodents (~23.7 hours for wild-type mice in constant darkness) versus humans (24.3-25.1 hours) (Kavakli & Sancar, 2002).

In addition to its circadian role, PER2 exhibits tumor suppressive functions as a transcriptional suppressor in many cancers (Fu, Pelicano, Liu, Huang, & Lee, 2002; Hwang-Verslues et al., 2013; Papagiannakopoulos et al., 2016). Suppression of PER2 at both gene expression and protein levels facilitates tumor growth, invasion and cancer malignancy (Chen et al., 2005; Hwang-Verslues et al., 2013; Winter, Bosnoyan-Collins, Pinnaduwage, & Andrulis, 2007; Yu et al., 2018). Since PER2 protein level oscillates and PER2 shuttles between nucleus and cytoplasm, its protein stability and subcellular localization may also determine its tumor suppression function. Importantly, SUMO pathways are often dysregulated in cancers (Seeler & Dejean, 2017). Such dysregulation could alter PER2-SUMO conjugation and subsequently inhibit PER2 tumor suppression function by either facilitating its degradation or inhibiting its nuclear entry. Thus, SUMO1 versus SUMO2 conjugation forms a critical crossroads controlling PER2 fate and influencing PER2 activities which extend beyond core circadian regulation. Discovery of this crossroads determining PER2 function provides new insights into the regulation of PER2 specifically as well as the function of SUMOylation in circadian regulation and health more broadly.

## Materials and Methods

### Plasmids and reagents

pcDNA3.1Myc-hPER2 plasmid was generously provided by Dr. Randal Tibbetts (University of Wisconsin, USA) (Shanware et al., 2011). pEGFP.C3.SUMO.GG plasmids were kindly provided by Dr. Mary Dasso (NIH, USA). pcDNA3.1.HA.Ub plasmid was kindly provided by Dr. Hsiu-Ming Shih (Academia Sinica, Taiwan). A series of mutant PER2 plasmids including S480A, S480D, K83R, K1163R, K736R, K83/1163R, K83/736R, K736/1163R, K83/736/1163R were made using site-directed mutagenesis (Agilent, Santa Clara, CA) with primer sets listed in Table S1. Depending on the experiments, these PER2 cDNAs were sub-cloned into various tagged vectors including pcDNA3.0.HA (Thermo Fisher Scientific, Waltham, MA), 3XFLAG-CMV-7.1-2 (Sigma-Aldrich, St. Louis, MO) and pcDNA3.1.Myc.HisA/pcDNA3.1.Myc.HisC (Thermo Fisher Scientific) for transient expression in mammalian cells. pGL3.PER2.Luc promoter (−948~+424) was generated as previously described (Yu et al., 2018). All cloned and mutated genes were verified by sequencing. The lentiviral shCtrl (TRC2.Scramble, ASN0000000003), shSUMO1 (TRCN0000147057), shSUMO2#1 (TRCN0000007653), shSUMO2#2 (TRCN0000007655), shRANBP2#1 (TRCN0000272800) and shRANBP2#2 (TRCN0000272801) were purchased from the National RNAi Core Facility (Taipei, Taiwan). Transient transfection was done by Mirus TransIT-LT1 Reagent (Mirus Bio, Madison, WI) according to the manufacturer’s instructions or by electroporation. Subcellular protein fractionation was performed using a kit for cultured cells from Thermo Fisher Scientific (78840). Proteasome inhibitors MG132, SUMO protease inhibitor N-Ethylmaleimide (NEM), cycloheximide and D-luciferin were from Sigma-Aldrich. CK1δ/ε inhibitor PF 670462 was purchased from TOCRIS (Minneapolis, MN).

### Cell line

Immortalized human bone osteosarcoma cell U2OS and embryonic kidney cell HEK-293Twere maintained in DMEM supplemented with 10% fetal bovine serum and antibiotics in a humidified 37 °C incubator supplemented with 5% CO_2_. Serum shock was performed with 50% horse serum (Thermo Fisher Scientific) for 2 h when cells reached ~90% confluence. The medium was then replaced with medium supplemented with 0.1% FBS. At the time indicated, the cells were washed twice with ice-cold PBS and whole-cell lysates prepared using RIPA lysis buffer (50 mM Tris-HCl (pH7.4), 150 mM NaCl, 1% NP40, 0.25% sodium deoxycholate, 0.05% SDS, 1 mM PMSF, 1x protease inhibitor, with or without 10 mM NEM).

### Generation of PER2^K736R^ knock-in mutant

An all-in-one CRISPR vector, pAll-Cas9.Ppuro (RNAi core facility, Academia Sinica, Taiwan), was digested with BsmBI and ligated with annealed oligonucleotides (GGCTGCACACACACAGAAGG) for sgRNA expression to target the PER2 coding region at the K736 residue. The 5’ mismatched G of PER2 sgRNA was created for optimal U6 transcriptional initiation. A single-stranded oligodeoxynucleotide (ssODN) (5‘-AAGGAGGTACTCGCTGCACACACACAG<u>CGC</u>GAGGAGCAGAGCTTCCTGCAGAAGTTCAAA-3’) was used to generate the K736R mutation. The underlined bases indicate the mutated codon encoding Arginine (R). Cells were transfected with all-in-one CRISPR plasmid and ssODNs donor template. After selection with puromycin, genomic DNA from selected cells was sequenced using a target-specific sequencing primer (5‘-GCACTAGCCTGGGACTATTC-3’) to validate the knock-in mutation.

### RNA isolation, reverse transcription and real-time (q)PCR

Total RNA from cell culture was isolated using Trizol reagent or RNeasy kit (Qiagen, Hilden, Germany) and reverse-transcribed with Quant iNova Reverse Transcription kit (Qiagen). qPCR was performed using Quant iNova SYBR Green PCR kit (Qiagen) and primer sets for gene expression and analyzed on a Applied Biosystems 7300 Real-Time PCR System (Thermo Fisher Scientific). Primers used in this study were listed in Table S2. Glyceraldehyde 3-phosphate dehydrogenase (GAPDH) mRNA was used as an internal control for mRNA expression. Expression levels were calculated according to the relative ΔCt method.

### Co-immunoprecipitation (Co-IP)

Whole-cell lysates were prepared using a TNE lysis buffer (10 mM Tris-Cl (pH 7.5), 150 mM NaCl, 0.5 mM EDTA, 1% NP-40, 10 μM MG132, 1 mM PMSF, 10 mM NEM and 1x protease inhibitors) followed by sonication and centrifugation at 12,000 × g at 4°C. 1 ml of the crude whole-cell extract was incubated with 1-2 μg of antibody as indicated or control IgG antibodies at 4°C overnight. Then, 50 μl prewashed protein A/G agarose was added to the mixture and incubated at 4°C for 4 h with gentle agitation. After extensive washing with diluted NP-40 lysis buffer (0.1-0.5% NP-40), PER2 interacting proteins were eluted with SDS buffer and analyzed by immunoblot. Antibodies and agarose used for PER2 IP were listed in Table S3.

### Denatured-IP

Denatured whole-cell lysates were prepared using a lysate buffer containing 1% SDS, 20 mM Tris-HCl (pH 8.0), 10% glycerol, 1 mM DTT, 15 mM NEM and 1x protease inhibitors, followed by boiling at 95°C for 8 minutes and sonication. After centrifugation at maximum speed, Flag-SUMO conjugated proteins were immunoprecipitated over night at 4°C. The samples were washed using high salt RIPA buffer (20 mM Tris-HCl (pH 8.0), 0.5 mM EDTA, 250 mM NaCl, 0.5% NP40, 10% glycerol, 10 mM NEM, 1xprotease inhibitors), low salt RIPA buffer (20 mM Tris-HCl (pH 8.0), 0.5 mM EDTA, 150 mM NaCl, 0.5% NP40, 10% glycerol, 10 mM NEM, 1xprotease inhibitors), and PBS before being boiled in 2xsample buffer (with 100 mM DTT) and processed for SDS-PAGE followed by immunoblot.

### in situ Proximity ligation assay (PLA)

U2OS cells were seeded at a density of 1.2×10^5^ cells on 12-mm glass cover slips (Hecht Assistent) 24 hours before serum shock. Cells were fixed in 4% paraformaldehyde for 10 min and then permeabilized in 4% paraformaldehyde containing 0.15% Triton for 15 min. Immunofluorescence staining was carried out using rabbit-anti-PER2 (1:250 dilution, Santa cruz, sc-25363) and mouse-anti-SUMO1 (1:350 dilution, Sigma, SAB1402954) antibodies. The corresponding Duolink® PLA Probes anti-rabbit PLA PLUS and anti-mouse PLA MINUS (1∶5 dilution) were used. Probe incubation, ligation and amplification reaction were performed using Duolink® PLA Reagents (Sigma) following the manufacturer’s instruction. Cells were examined with a Leica confocal SP8 microscope (objective× 63). For each cover slip, 9 non-overlapping images were taken. Cells with positive signals in nucleus only versus signals in both nucleus and cytoplasm were counted at each CT. At least two independent experiments were performed.

### Immunoblotting (IB)

Whole-cell lysates were prepared using RIPA lysis buffer (50 mM Tris-HCl (pH7.4), 150 mM NaCl, 1% NP40, 0.25% sodium deoxycholate, 0.05% SDS, 1 mM PMSF, 1x protease inhibitor, 10 mM NEM). IB analysis was performed after 6-7.5% SDS-PAGE or 4-20% Mini-PROTEAN TGX Precast Protein Gels (BioRad, Hercules, CA), with overnight incubation with a 1:1000 dilution of primary antibody and followed by a 1:5000 dilution of horseradish peroxidase-conjugated anti-rabbit or anti-mouse antibody (Jackson ImmunoResearch, West Grove, PA). Signals were detected using Millipore Immobilon Western Chemiluminescent HRP Substrate (Merk Darmstadt, Germany). Primary antibodies used in this study are listed in Table S3. Protein concentration was determined by the Bradford assay (Bio-Rad) before loading and verified by α-tubulin level. The optical density was determined using the National Institutes of Health ImageJ program.

### Nano LC-MS/MS Analysis

In order to identify the sumoylation site(s) of PER2, flag-PER2 which was expressed with and without over-expressed SUMO1(RGG) were purified by SDS-PAGE and followed by in-gel digestion for the mass spectrometric analysis. Briefly, gel slices were cut into 1 mm squares and washed in 50 mM triethylammonium bicarbonate buffer (TEABC) and then TEABC/ACN (25 mM TEABC, 50% acetonitrile). Repeated the washing steps three times. The gel slices were reduced in 20 mM dithiothreitol (DTT) at 56 °C for 1 h, and alkylated in 55 mM iodoacetamide at room temperature for 45 min in dark before trypsin digestion at 37 °C overnight. The enzymatic digestions were quenched through the addition of formic acid (10%) and the tryptic peptides were collected and vacuum-dried prior to mass analysis.

MS data were acquired on an Orbitrap Fusion mass spectrometer equipped with Dionex Ultimate 3000 RSLC system (Thermo Fisher Scientific) and nanoelectrospry ion source (New Objective, Inc., Woburn, MA). Samples in 0.1% formic acid were injected onto a self-packed precolumn (150 μm I.D. × 30 mm, 5 μm, 200 Å) at the flow rate of 10 μl min-1. Chromatographic separation was performed on a self-packed reversed phase C18 nano-column (75 μm I.D. x 200 mm, 3 μm, 100 Å) using 0.1% formic acid in water as mobile phase A and 0.1% formic acid in 80% acetonitrile as mobile phase B operated at the flow rate of 300 nL min-1. The MS/MS were run in top speed mode with 3s cycles; while the dynamic exclusion duration was set to 60 s with a 25 ppm tolerance around the selected precursor and its isotopes. Monoisotopic precursor ion selection was enabled and 1+ charge state ions were rejected for MS/MS. The MS/MS analyses were carried out with the collision induced dissociation (CID) mode. Full MS survey scans from m/z 300 to 1,600 were acquired at a resolution of 120,000 using EASY-IC as lock mass for internal calibration. The scan range of MS/MS spectra based on CID fragmentation method were from m/z 150 to 1,600. The precursor ion isolation was performed with mass selecting quadrupole, and the isolation window was set to m/z 2.0. The automatic gain control (AGC) target values for full MS survey and MS/MS scans were set to 2e6 and 5e4, respectively. The maximum injection time for all spectra acquisition was 200 ms. For CID, the normalization collision energy (NCE) was set to 30%.

For data analysis, all MS/MS spectra were converted as mgf format from experiment RAW file by msConvert then analyzed by Mascot for MS/MS ion search. The search parameters included the error tolerance of precursor ions and the MS/MS fragment ions in spectra were 10 ppm and 0.6 Da and the enzyme was assigned to be trypsin with the miss cleavage number two. The variable post-translational modifications in search parameter were assigned to include the oxidation of methionine, carbamidomethylation of cysteine, and GlyGly tag of lysine as the sumoylation site.

### Transient reporter assay

HEK-293T cells (1.2×10^5^ cells) without or with SUMO1 depletion were transfected with 20 ng pGL3.PER2.Luc promoter, 5 ng Renilla.Luc (for normalization) and FLAG-PER2^WT^ (0, 20, 40 ng) plasmids in a well of a 12-well plate. Cell extracts were prepared at 24 h after transfection, and the luciferase activity was measured using the Dual-Glo Luciferase Assay System (Promega).

### Bioluminescent recording and data analysis

CRISPR/CAS edited U2OS cells (4×10^5^ cells) were seeded in a 35-mm plate a day before transfection with 1 μg PER2.Luc reporter. Twenty hours after transfection, the cells were reached ~90% confluent and synchronized with 100 nM dexamethasone (Balsalobre et al., 2000) for 2 h. Subsequently, the media was changed to recording media (Gibco phenol red free DMEM (Thermo Fisher Scientific), 400 μM D‐Luciferin Potassium Salt (Goldbio, St Louis MO) and monitored via the use of a Kronos Dio luminometer (AB-2550, ATTO, Tokyo, Japan) at 37°C. Bioluminescence from each plate was continuously recorded (integrated signal of 60 s with intervals of 9 min). Data were smoothed and detrended with the KronosDio analysis tool. JTK_Cycle analysis (version 3) was employed to test the rhythmicity and amplitude of the signals between circadian time 5 and 31 hours (Hughes, Hogenesch, & Kornacker, 2010; M. Miyazaki et al., 2011). The phase shift was determined by measuring the time difference between the peaks of the PER2 promoter luciferase rhythms between the PER2^WT^ and PER2^K736R^ cells.

### Statistical analysis

All data are presented as means ± SD. Student’s *t*-test was used to identify statistically significant differences. Asterisk (*) indicates statistical significance with *p*-value <0.05, (**) indicates *p*-value < 0.01, and (***) indicates *p*-value < 0.001.

## Acknowledgements

This work was supported by Academia Sinica and Ministry of Science and Technology [MOST 105-2628-B-001-008-MY3] and [MOST-108-2311-B-001-005-MY3] to WWHV. We thank the National RNAi Core Facility (Academia Sinica) for RNAi and CRISPR reagents. We also thank Mr. Kuo-Chih Cheng and Dr. Ming-Ta Lee for their assistance and discussion.

## Author contributions

Conceptualization, L.C.C., Y.L.H. and W.W.H.-V; Methodology, L.C.C., Y.L.H., Y.C.C., P.H.H., W.W.H.-V.; Investigation, L.C.C., T.Y.K.; Writing – Original Draft, L.C.C. and W.W.H.-V.; Writing – Review & Editing, L.C.C., Y.L.H. and W.W.H.-V.; Funding Acquisition, W.W.H.-V.; Supervision, W.W.H.-V.

## Supplementary Figures and Tables

**Figure S1.**
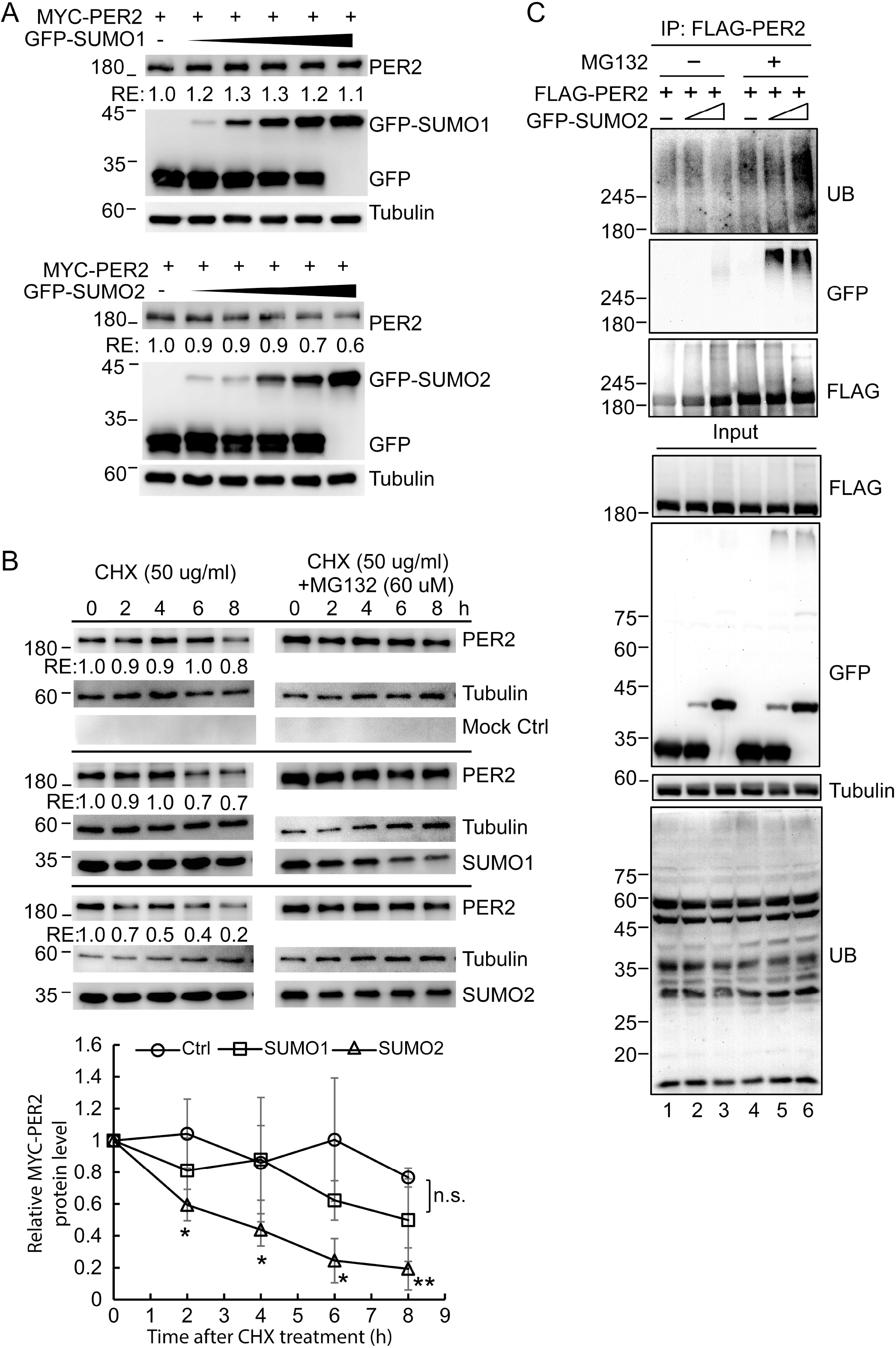
SUMO2 conjugation promotes PER2 protein ubiquitination and degradation. **A.** IB assay of PER2 and SUMOs using U2OS cells co-transfected with MYC-PER2 and increasing amount of GFP-SUMOs. Tubulin was used as a loading control. RE: relative expression. Relative MYC-PER2 protein levels from three IB analyses were quantified. Data are means ± SD. **B.** Top: Time course assay using cycloheximide (CHX) treated HEK-293T cells co-transfected with MYC-PER2 and GFP-SUMOs. Cells were treated without or with MG132. The levels of PER2 and SUMOs were determined using IB analysis. Tubulin was used as a loading control. RE: relative expression. Bottom: Quantification of relative PER2 protein levels from three IB analyses. Data are means ± SD. n.s., non-significant; *, *p* < 0.05 (Student’s t-test). **C.** Co-IP assay using HEK-293T cells co-transfected with FLAG-PER2 and GFP-SUMO2. Cells were treated without or with proteasome inhibitor MG132. Cell lysates were IP with anti-FLAG antibody and analyzed by IB to detect PER2-SUMO and −UB conjugates. The experiment was performed twice with similar results.

**Figure S2.**
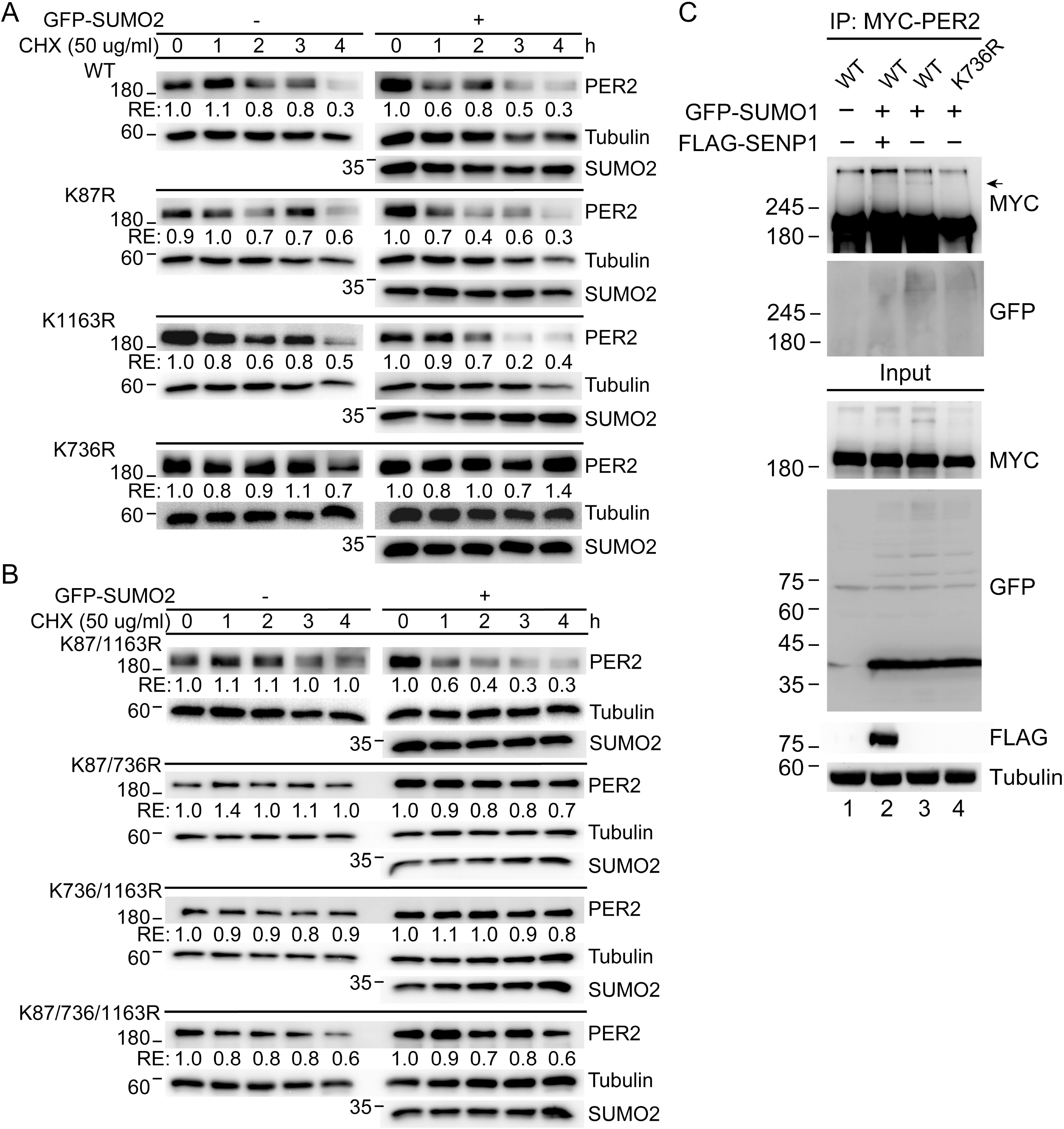
Lysine residue K736 is a critical site for PER2 SUMOylation. A. Time course assay using CHX treated HEK-293T cells co-transfected with K-to-R single-site mutated MYC-tagged PER2 with or without GFP-tagged SUMO2. The levels of PER2 and SUMO2 were determined using immunoblotting. Tubulin was used as a loading controls. RE: relative expression. B. Time course assays using CHX treated HEK-293T cells co-transfected with K-to-R double- or triple-site mutated MYC-tagged PER2 with or without GFP-tagged SUMO2. The levels of PER2 and SUMO2 were determined using immunoblotting. Tubulin was used as a loading controls. RE: relative expression. C. Co-IP assay using HEK-293T cells co-transfected with MYC-PER2, GFP-SUMOs and FLAG-SENP1. Cell lysates were IP with anti-MYC antibody. Immunoprecipitates and input lysates were analyzed by IB. Arrow indicates the PER2-SUMO conjugates. All experiments were performed at least twice with similar results.

**Figure S3.**
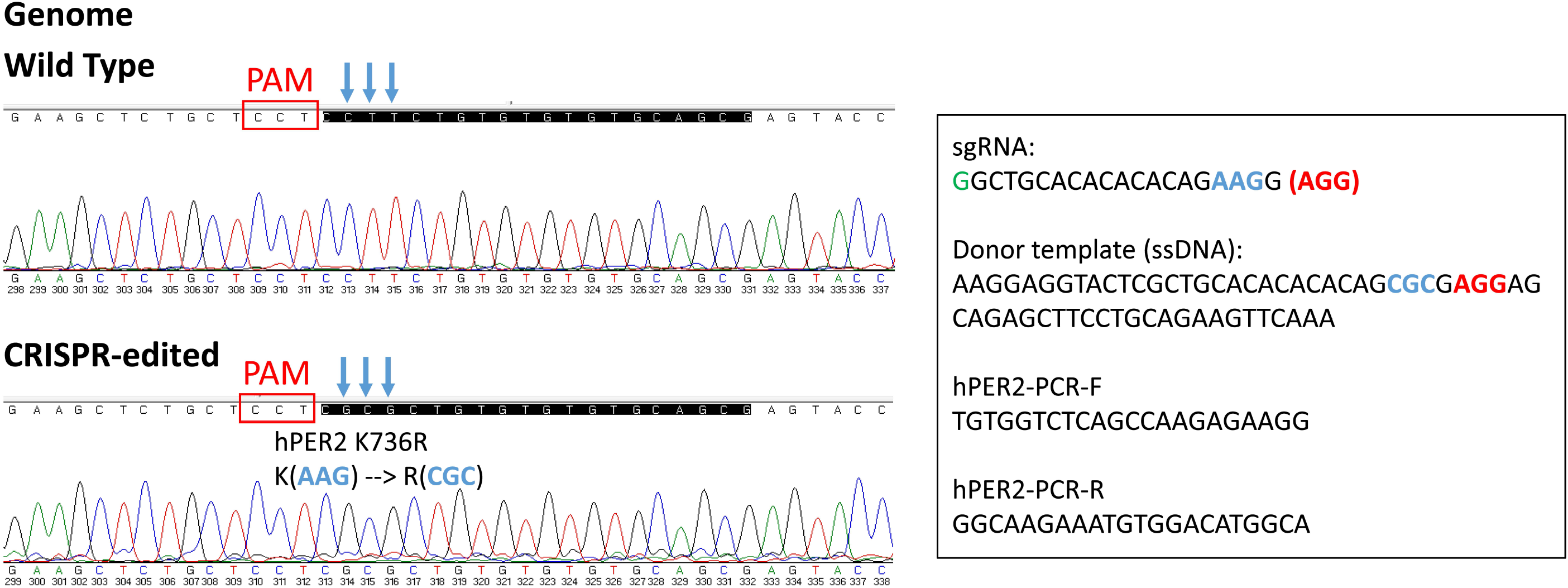
Generation of PER2^K736R^ knock-in mutant HEK-293T cells. The WT and K736R PER2 sequences, 5’-mismatched G of PER2 sgRNA, donor template and PCR primers are shown.

**Figure S4.**
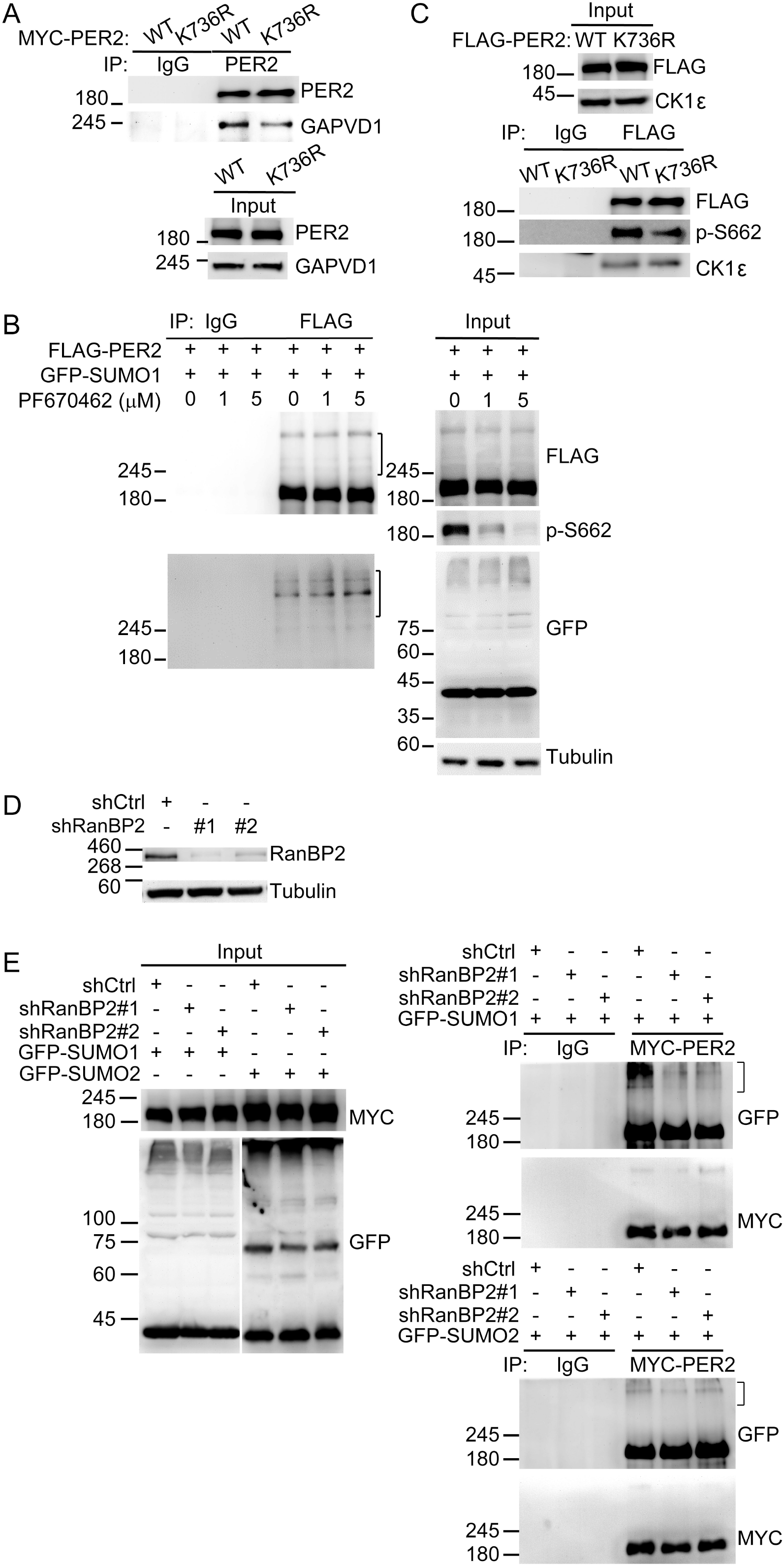
PER2 K736-dependent SUMO1 conjugation promotes CK1 phosphorylation of PER2 S662 and CRY1/GAPVD1 interactions required for PER2 nuclear translocation and PER2-mediated transcriptional suppression. **A.** Co-IP assay of PER2 and GAPVD1 using U2OS cells transfected with MYC-PER2^WT^ or −PER2^K736R^. Cell lysates were IP with indicated antibodies and analyzed by IB. Normal IgG was used as a control. The experiment was repeated three times. **B.** Co-IP assay using HEK-293T cells co-transfected with FLAG-PER2^WT^ and GFP-SUMO1 treated without or with CK1 inhibitor PF670462. Cell lysates were IP with anti-FLAG antibody and analyzed by IB to detect PER2-SUMO conjugates. Normal IgG was used as an IP control. The inhibition efficiency of CK1 inhibitor was determined using IB analysis for PER2 S662-phosphorylation. **C.** Co-IP assay using U2OS cells co-transfected with FLAG-PER2^WT^ or −PER2^K736R^. Cell lysates were IP with anti-FLAG antibody and analyzed by IB to detect CK1 and S662-phosphorylated PER2. **D, E.** The depletion efficiency of RanBP2 protein was determined using IB analysis (D). Co-IP assay using HEK-293T cells without or with RanBP2 depletion co-transfected with MYC-PER2 and GFP-SUMOs (E). Cell lysates were IP with anti-MYC antibody and analyzed by IB to detect PER2-SUMO1 (E, upper right panel) and PER2-SUMO2 (E, lower right panel) conjugates indicated by brackets. All experiments were performed at least twice with similar results.

**Figure S5.**
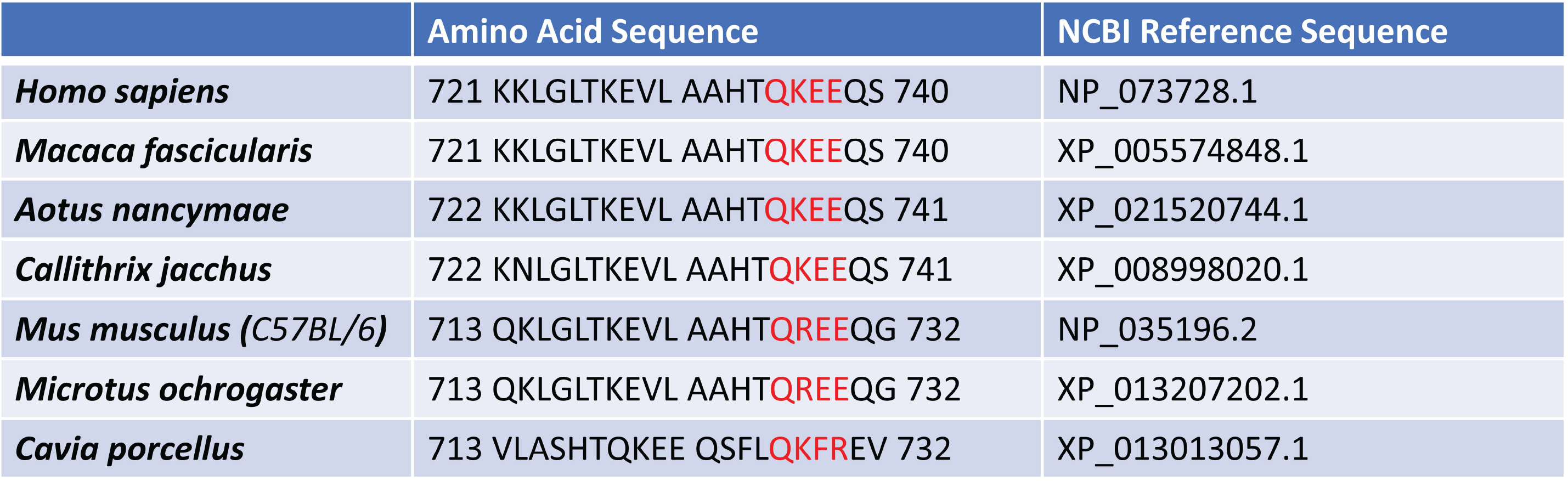
PER2 protein sequence alignment of the region containing human K736 (QKEE motif) between primates and rodent.

**Table S1.**
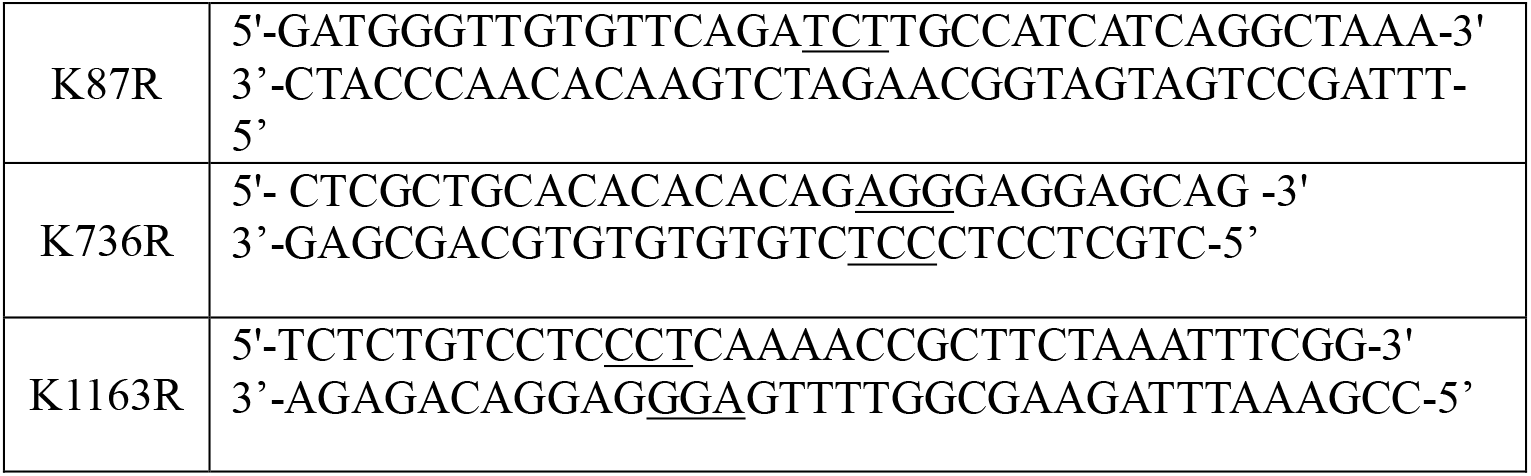
Primers for site-directed mutagenesis.

**Table S2.**
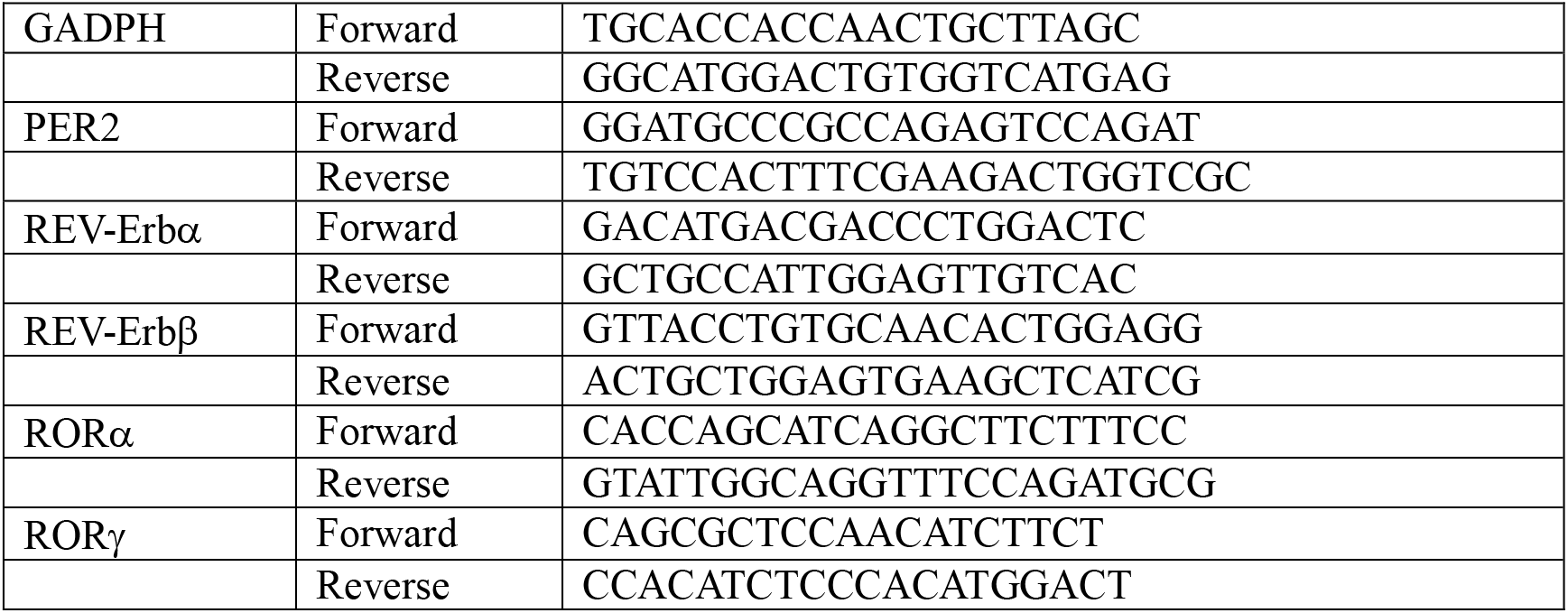
Primer sets for qPCR.

**Table S3.** Antibodies.

| **Antibody** | **Catalog number** | **Dilution** | **Company** |
| --- | --- | --- | --- |
| CRY1 | SC-101006 | 1:1000 | Santa Cruze |
| CK1ε | 12448 | 1:1000 | Cell signaling |
| GFP | 2555S | 1:1000 | Cell signaling |
| GAPex5 (GAPVD1) | A302-116A | 1:1000 | Bethyl |
| PER2(H90) | SC-25363 | 1:1000 | Santa Cruze |
| PER2 (phosphor S662) | Ab206377 | 1:1000 | Abcam |
| RanBP2 | Ab64276 | 1:1000 | Abcam |
| SUMO1 | PA5-34765 | 1:1000 | Invitrogen |
| SUMO2 | PA5-11373 | 1:1000 | Thermo |
| β-TrCP | 4394S | 1:1000 | Cell signaling |
| UB | ab7780 | 1:1000 | Abcam |
| HA.11 epitope Tag | 901503 | 1:1000 | BioLegend |
| FLAG-M2 | F3165 | 1:1000 | Sigma |
| Myc-tag | ab9106 | 1:1000 | Abcam |
| alpha tubulin | GTX628802 | 1:10000 | GeneTex |
| Nuclear Matrix Protein p84 | GTX70220 | 1:1000 | GeneTex |

For Co-IP assays, 1-2 ug per IP was used.

